# Chaperone saturation mediates translation and protein folding efficiency

**DOI:** 10.1101/2025.06.25.661590

**Authors:** Andrew T. Martens, Vincent J. Hilser

## Abstract

Whether the emergence of a nascent protein from the ribosome and the formation of structural elements are synchronized has been a longstanding question (Chaney and Clark, 2015; Deane and Saunders, 2011; Levinthal, 1968; Marin, 2008; Sauna and Kimchi-Sarfaty, 2011; Spencer and Barral, 2012; Tsai et al., 2008; Zhang and Ignatova, 2011). Paradoxically, kinetically efficient translation can induce mis-folding and aggregation despite the presence of molecular chaperones (Siller et al., 2010; Spencer et al., 2012), which in *Escherichia coli* are induced by unfolded protein (Parsell and Sauer, 1989) *via* σ^32^ (Craig and Gross, 1991). The molecular mechanisms mediating translation efficiency and protein folding efficiency remain poorly understood. Using ribosome profiling (Ingolia et al., 2009) and protein quantitation, we show that synonymous changes to Firefly *Luciferase* (*Luc*) mRNA have a direct effect on its translation efficiency. These changes alone cause up to a 70-fold difference in Luc protein levels. However, increased Luc protein is met with at most a ∼2-fold increase in chaperone levels, revealing that the σ^32^ transcriptional response has saturable properties. This response is found to be poised near its midpoint (where it is most sensitive to perturbation) when *Luc* mRNA has an intermediate translation efficiency. These results suggest not only that chaperone saturation limits the ability of cells to maintain protein folding homeostasis when challenged with highly efficient translation, but that translation efficiency and protein folding efficiency evolved for mutual sensitivity.

## Results

Ribosome profiling provides two related types of information: the relative statistics for ribosome occupancy at each codon position in a given mRNA, which should relate to the local translation elongation rate (Martens et al., 2015), and the allocation of ribosomes to each mRNA as a fraction of all ribosomes, which relates to the amount of protein produced (Li et al., 2014). To resolve how kinetically efficient *Luc* translation causes Luc protein folding defects, we performed ribosome profiling on *E. coli* expressing synonymously re-coded *Luc* mRNA variants that were modified to change the translation efficiency, but not the amino acid sequence (**Table S1**) (**Methods** and **Supplementary Information**). This test set contains several variants which have been demonstrated to have different aggregation propensities and experimentally determined translation rates (Spencer et al., 2012), as well as 4 new hybrid *Luc* constructs, each of which contains a transition between high and low translation efficiency regimes. Two constructs contain transitions (from high to low, and low to high) at codon 16, near the start codon, while two other constructs contain a transition at codon 293, in the center of the sequence. If these transitions decrease or increase the efficiency of translation, they should decrease or increase, respectively, the spacing between consecutive ribosomes, and cause corresponding increases or decreases, respectively, in average ribosome occupancy along the mRNA. The test set also includes two “wild-type” constructs, one with the native codons as found in the firefly mRNA (Luc WT*_P. pyralis_*), and the other re-coded to match the translation efficiency from the firefly into *E. coli* (Luc WT*_E. coli_*) (Spencer et al., 2012).

Comparison of the ribosome profiles for the different variants reveals two significant features. First, a ribosome occupancy peak is evident within the first ∼50 codons (**Figure 1**), a result that is consistent with ribosome pausing as the nascent protein initially emerges from the ribosome (Oh et al., 2011). Second, efficient translation is associated with low ribosome occupancies, while inefficient translation is associated with high ribosome occupancies. Ribosome occupancy gradually increases towards the 5ʹ end along S_2-550_ mRNA, but is more evenly distributed across the F_2-550_ mRNA (**Figure 1 a&b**). This difference is more pronounced for S_2-16_F_17-550_ and F_2-16_S_550_ mRNA (**Figure 1 c&d**). Conversely, ribosome occupancy sharply increases, then decreases, for S_2-292_F_293-550_ mRNA, and continually increases for F_2-292_S_293-550_ mRNA (**Figure 1 e&f)**. In comparison, the two wild-type sequences have relatively more uniform ribosome occupancy profiles (**Figure 1 g&h**). Synonymously re-coding the Luc mRNA therefore changes its ribosome profile, consistent with standard ribosome queuing models which relate ribosome occupancy profiles to the flux of ribosomes along the mRNA (von der Haar, 2012).

**Figure 1:**
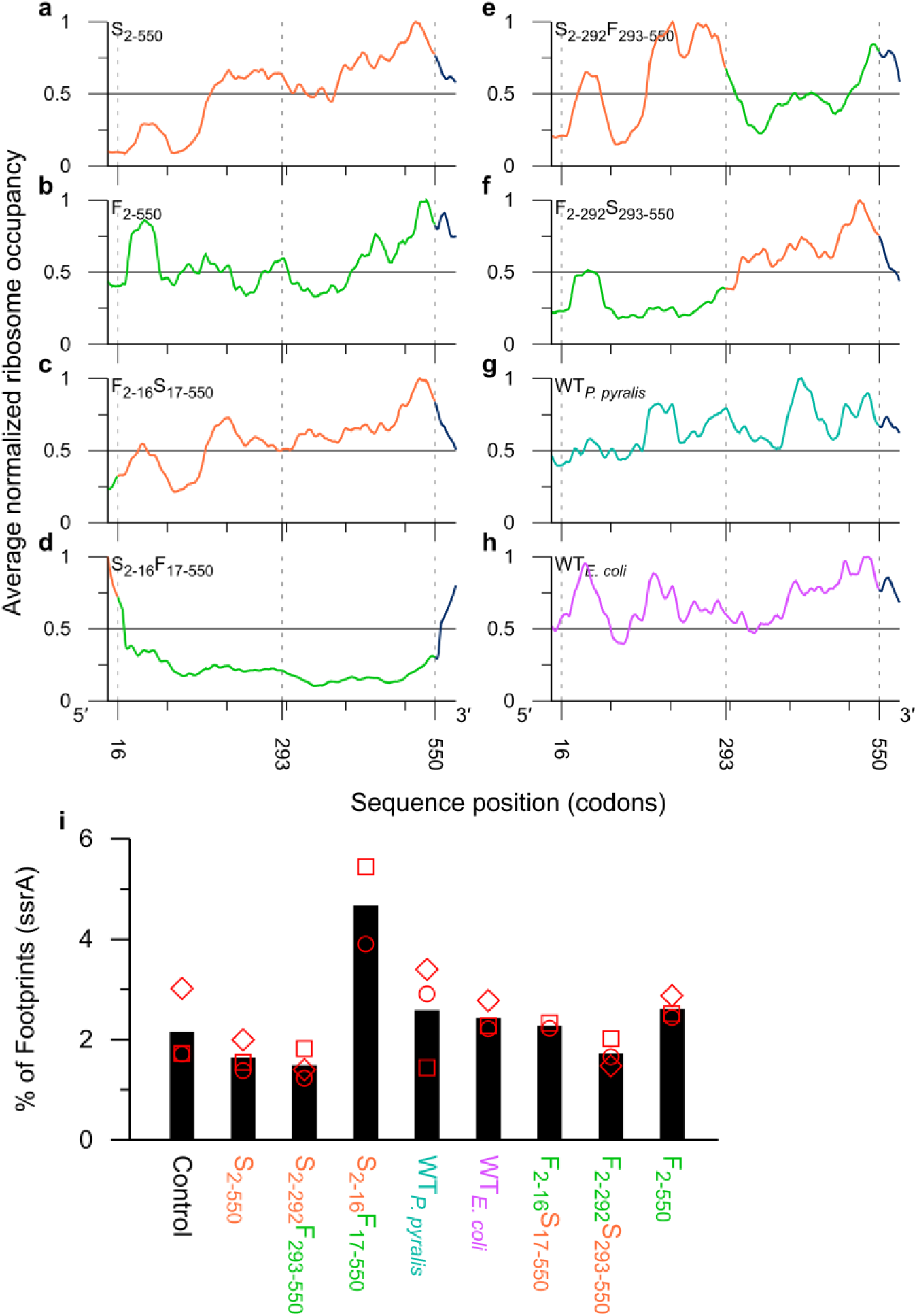
The mRNA sequence determines the *Luc* ribosome profile and ribosome spacing. **a-h:** Measured ribosome occupancy effects of wild-type, slow (orange) and fast codons (green) at positions 2 – 16, 17 – 292, and 293 – 550. The C-terminal tag was not re-coded (blue, positions 551 – 584). Ribosome occupancy was normalized from 0 to 1 and averaged across replicates (n = 2 or 3). **i:** Relative abundance of ribosome-protected tmRNA as a function of *Luc* mRNA sequence. Symbols: experimental replicates (n = 2 or 3). Bars: means of the replicates.

As a further test for whether recoding changes ribosome occupancy, we analyzed the ssrA-mediated mRNA degradation pathway. Because ribosome occupancy changes as a function of the mRNA translation efficiency, it follows that the average spacing between sequential ribosomes is reduced on mRNAs with high ribosome occupancy and increased on mRNAs with low occupancy. Indeed, a connection has been suggested between ribosome spacing and mRNA degradation (Hanson and Coller, 2018; Kennell, 2002), with greater spacing between ribosomes exposing the mRNA to endonucleolytic attack, generating a truncated mRNA without a stop codon. In *E. coli*, stalled ribosomes are rescued in a process which involves recruitment of tmRNA, a species which is ribosome-protected and can therefore be directly detected using ribosome profiling (Guydosh and Green, 2014), followed by degradation of the nascent polypeptide after tagging with the ssrA peptide (Keiler, 2015).We therefore predict, in accordance with ribosome queuing models, activation of this system in cells with increased spacing between ribosomes translating *Luc* mRNA. As expected, only the Luc S_2-16_F_17-550_ sample, which combines poor translation efficiency at the start of the mRNA with high translation efficiency and rapid clearing of ribosomes across the remainder of the mRNA, shows an increase in the abundance of ribosome-protected tmRNA (**Figure 1 i**). Taken together, the differences in ribosome occupancy data for the different variants and the tmRNA effect provide strong evidence that Luc translation efficiency and ribosome occupancy are predictably changed as a function of the *Luc* mRNA sequence.

To determine how the differences in translation efficiency are related to Luc protein production, folding efficiency and chaperone synthesis, we again turned to ribosome profiling. First, we interrogated the same ribosome profiling data to determine the percent allocation of ribosomes (Li et al., 2014) to the *Luc* mRNA. We find that the *Luc* mRNA sequence has a strong effect on its ribosome allocation, and the identity of the first sixteen codons is of particular importance (**Figure 2 a**), consistent with other studies on the effects of codons near the start codon (Firnberg et al., 2014; Kudla et al., 2009; Pop et al., 2014). We infer that efficient translation near the start codon results in more rapid vacating of the start site and subsequent initiation events, leading to high allocation levels. Conversely, inefficient translation around the start site leads to low allocation levels. To determine how ribosome allocation is related to final protein production, we monitored total Luc protein using immunoblotting, and functional Luc protein using enzymatic activity (**Figure 2 b&c**). Both total and active Luc protein are highly correlated to each other (r^2^ = 0.99, **Figure S1**) (with the notable exception of Luc S_2-16_F_17-550_), confirming that the efficiency of translation immediately downstream of the start codon plays a major role in determining overall Luc protein levels, and that Luc protein synthesis rates are well within the regime suitable for making active protein. Interestingly, we also note that total protein levels are greatest for construct F_2-550_, although this construct has relatively low ribosome allocation compared to F_2-16_S_17-550_ and F_2-292_S_293-550_. This effect is explained by efficient transit of ribosomes along the entire mRNA, resulting in more proteins produced per mRNA over time and fewer ribosomes bound. An opposite effect is also observed for construct S_2-16_F_17-550_. Of note is that translation of the endogenous *lacZ* mRNA, whose gene has the identical promoter and RBS sequences as the *Luc* genes and is induced along with *Luc*, is unchanged (**Figure 2 a**) indicating that the effects are not due to changes in the cell availability of ribosomes but are due to the interactions of the specific Luc mRNA of each variant with a uniform concentration of ribosome.

**Figure 2:**
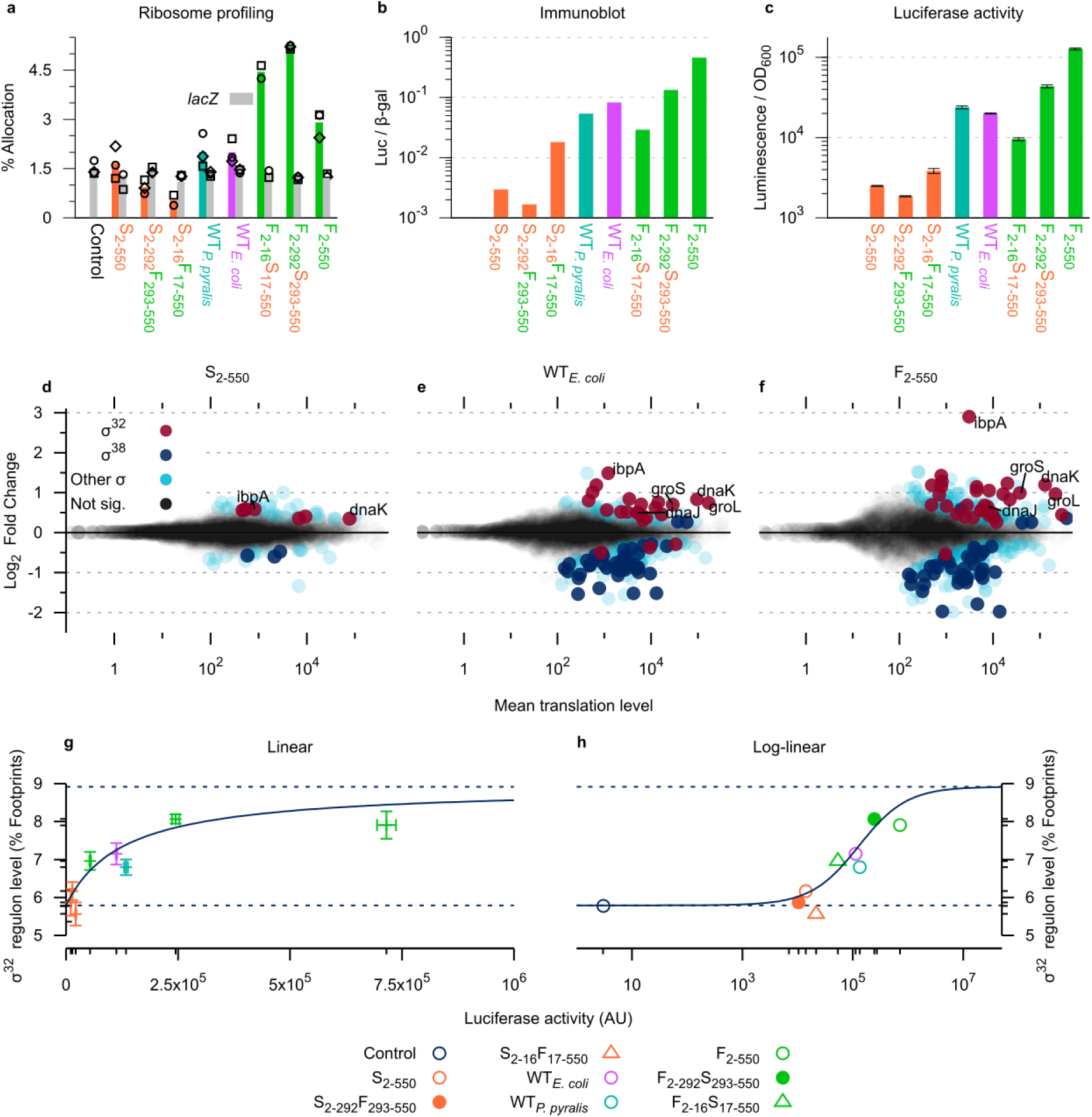
Activation of σ^32^ depends on *Luc* mRNA sequence and Luc protein levels, and exhibits saturable properties. **a.** Quantification of ribosome-protected *Luc* (color) and *lacZ* (grey) mRNA fragments (symbols: replicates; n = 2 or 3). **b.** Quantification of Luc protein relative to β-galactosidase protein by immunoblotting (arbitrary units). **c.** Quantification of Luc enzymatic activity (arbitrary units; bars: 95% CI; n = 3). **d – f.** Fold change (relative to Luc negative cells) and mean translation levels for the *E. coli* transcriptome as a function of the *Luc* mRNA sequence (**d:** S_2-550_, **e:** WT*_E. coli_*, **f:** F_2-550_). Red and blue circles: σ^32^ regulon and σ^38^ regulon, significant (p_adj_ < 0.10). Light blue circles: significant, other regulons. Grey circles: not significant. **g & h.** Fit between Luc enzymatic activity and σ^32^ induction (**d:** linear, **e:** log-linear). Fit (blue, solid line) shown with the estimated basal and predicted maximal σ^32^ activities (blue dashed lines). Error bars: standard error of the mean using Orear’s effective variance method.

Because Luc protein requires chaperones to fold efficiently in *E. coli* (Agashe et al., 2004), translation of *Luc* mRNA stimulates chaperone synthesis *via* sigma factor σ^32^. As such, expression of Luc mRNAs with different sequences and translation efficiencies should differentially activate σ^32^. To test this hypothesis, ribosome profiling was used to directly monitor the cellular response to the production of Luc protein by quantifying translation of all transcripts which comprise the σ^32^ regulon. The results reveal a strong, positive relationship between the *Luc* mRNA sequence, Luc protein production and σ^32^ regulon translation (**Figure 2 d-f**). Individually, we note a significant increase in the translation of σ^32^ regulon mRNAs, including those coding for the major chaperones DnaK, DnaJ, GroEL, and GroES (**Figure 2 d-f**). Surprisingly, the most highly up-regulated gene encodes IbpA (Inclusion Body Binding Protein A), revealing an enhanced ability to process protein aggregates (LeThanh et al., 2005) in response to elevated Luc protein levels. This result, consistent with the existence of additional regulators of *ibpA* besides σ^32^ ^(Gaubig et al., 2011)^, may indicate that *E. coli* adapts to especially elevated levels of protein folding stress by providing for the ability to sequester misfolded proteins, which may otherwise exceed the saturation point of the chaperone load in the cell.

Globally, the first sixteen codons are most predictive of the relationship between the *Luc* mRNA sequence, Luc protein levels and σ^32^ activity (**Figure 2 d-f** and **Figure S2**), demonstrating that Luc protein activates σ^32^ in a concentration-dependent manner. To determine the total σ^32^ response level to Luc protein synthesis, total translation levels of the σ^32^ regulon were analyzed as a function of total Luc protein levels (**Figure 2 g&h**). Of note, we find σ^32^ activation in response to Luc protein to be non-linear, with an activity transition reminiscent of a binding equilibrium (**see Methods**). We also note that the midpoint of this transition, often indicative of the regime of greatest sensitivity in enzyme kinetics (Fersht, 1974), is reached when expressing the two wild-type constructs, suggesting the σ ^32^ mechanism is tuned to respond to surprisingly modest amounts of protein overproduction. Indeed, as would be expected by the sigma factor binding model, we observe a diminishing return of the σ^32^ response to the Luc stimulus (**Figure 2**), thus exhibiting saturable behavior. To validate the model and to determine whether the increase in chaperone mRNA is being driven by Luc-dependent sequestration of chaperones from σ^32^, we also measured the amount of translation from the σ^38^ regulon. As expected, activation of σ^32^ is accompanied by suppression of σ^38^, directly supporting of the sigma factor competition model (Maeda et al., 2000) (**Figure 2 a-c, dark blue circles**). Similarly, translation of trigger factor (0.4%; not regulated by σ^32^) (**Figure S3**) and of ribosomal proteins (14.5% (Dennis and Bremer, 1974; Schleif, 1967); regulated by σ^70^) (**Figure S4**) remain unaffected, further demonstrating that the effect is mediated specifically through σ^32^, which is controlled through its interactions with chaperones.

## Discussion

We have characterized the systemic impact of synonymous changes to the *Luc* mRNA by combining ribosome profiling with protein quantitation. These experiments make evident the major role that the *Luc* mRNA sequence plays in determining translation efficiency and ribosome occupancy. We also demonstrate that changes to *Luc* translation rates have predictable effects on Luc protein levels, with the greatest effect localized near the start codon. Most importantly, by comparing Luc protein levels to chaperone production levels, we find that translation rates, protein levels and chaperone availability are part of an integrated protein folding homeostatic system. The connections between these three aspects of protein regulation have broad implications in understanding several tradeoffs in protein production and folding.

The bacterial protein folding system may be considered in terms of DnaK/DnaJ (homologous to eukaryotic Hsp70/Hsp40) protein binding sites. During protein synthesis, unfolded or partially folded amino acid residues, belonging to the nascent protein, emerge from the ribosome into the aqueous cytosol (Lu and Deutsch, 2005). Exposed patches of hydrophobic residues may then bind DnaK/DnaJ or trigger factor co- or post-translationally (Deuerling et al., 1999; Teter et al., 1999). The σ^32^ auto-regulatory mechanism takes advantage of this phenomenon: the binding of DnaK to these hydrophobic patches precludes its binding to σ^32^, freeing σ^32^ to bind RNA polymerase and to activate the transcription of new chaperone mRNAs (Craig and Gross, 1991).

In the limit that the fraction of σ^32^ molecules bound to RNA polymerase approaches 1, one would predict diminishing returns when challenged with additional unfolded protein, resulting in saturation of the response and reduced protein folding efficiency. Our hypothesis finds support from numerous cases in the literature. For example, high production of proteins which depend on chaperones, or genetic ablation of crucial protein folding machinery of the cell, additively reduce the yield of folded protein (Agashe et al., 2004). Likewise, significant fractions of the MBP, GFP and Luc protein were insoluble when efficiently translated (Siller et al., 2010; Spencer et al., 2012), and the folding of bovine gamma-B crystallin was altered by synonymously changing its coding sequence in a trigger factor null genetic background (Buhr et al., 2016).

Other mechanisms regulating translation may also lead to chaperone saturation effects. Ribosome pauses could lead to depletion of chaperones or trigger factor by sequestering them to nascent proteins, remaining bound until translation is completed (Kaiser et al., 2006). It was shown that ribosome stalling at a pseudoknot structure activates σ^32^ (Gorochowski et al., 2019), consistent with DnaK binding co-translationally to the nascent proteins on the stalled ribosomes. Similarly, less efficient translation could shift the equilibrium between folded states (Sander et al., 2014) by promoting or inhibiting co- or post-translational DnaK or trigger factor binding. Our model may also extend to eukaryotic systems. For example, specific pause points in the mRNA encoding the eukaryotic P-gp (MDR1) protein (Kimchi-Sarfaty et al., 2007) might modulate co-translational chaperone binding and similarly deplete chaperones in the cell. Intriguingly, sequestration of chaperones to nascent proteins would be amplified the longer it takes to synthesize the nascent chain and release it from the ribosome, raising the possibility of a tradeoff between efficient chaperone binding and slower translation. In summary, the encoding of such co-translational schemes, in the context of a saturable chaperone response, would be highly sensitive to protein expression levels, mediating protein folding homeostasis in the cell.

Taken together, a growing body of evidence points towards to a unifying principle under which two opposing mechanisms, the synthesis of chaperones and of chaperone-dependent proteins, govern protein folding efficiency in the cell. The principle of chaperone saturation can be further extended to include any cellular stressors that affect the proteome, such as elevated temperature or denaturing agents. Thus, rather than attributing the folding efficiency of the native protein only to the rate of translation of its mRNA, our results suggest that folding efficiency is modulated as a function of the steady-state availability of chaperones to engage all proteins with exposed hydrophobic residues, either co- or post-translationally. This principle establishes the basis for a physical model linking protein folding homeostasis to any mechanism affecting translation rates (Leininger et al., 2019; Ninio, 2006).

There are several important implications of this result. First, we can now begin to quantitatively define the proteostatic limits for natural systems in terms of chaperone availability, and to gain insight into the tradeoffs between efficient protein production and faithful protein folding. Second, a deeper understanding of these tradeoffs will enable us to better engineer and express synthetic or foreign proteins while balancing protein levels with protein folding efficiency. Finally, we resolve the longstanding question of how synonymous sequences differentially affect protein folding by invoking canonical biological pathways in a new way.

## Acknowledgments

We thank José Barral, Allen Buskirk, Rachel Green, Nick Ingolia, Christian Kaiser, John Kim, Elijah Roberts, Robert Schleif and members of the Hilser lab for their advice, discussions and comments.

Funding from NIH (R01-GM063747 to V.J.H., R01-GM121567 to V.J.H., Cellular and Molecular Biology graduate student training grant 2T32-GM007231 to A.T.M.) and Johns Hopkins University is gratefully acknowledged.

## Author Contributions

A.T.M. and V.J.H designed the experiments. A.T.M. performed the experiments and analyzed the data. A.T.M. and V.J.H. wrote the manuscript.

## Supplementary Figures

**Figure S1:**
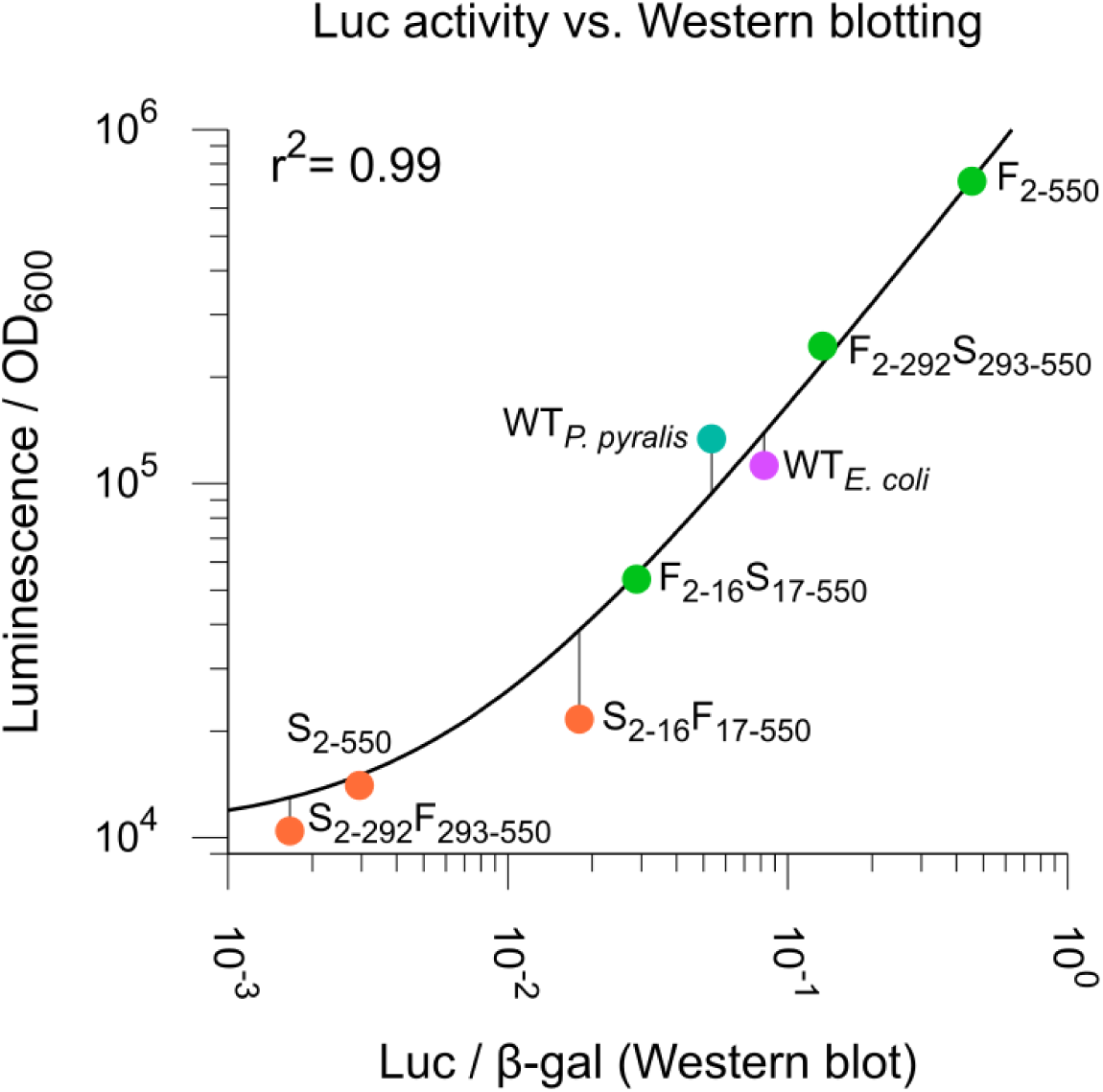
Measurements of Luc protein levels using enzyme activity assays (vertical axis) and immunoblotting (horizontal axis). Correlation coefficient (r^2^ = 0.99) was calculated after excluding S_2-16_F_17-550_ (statistical outlier, see main text). The linear fit is displayed along the points on a log-log plot.

**Figure S2:**
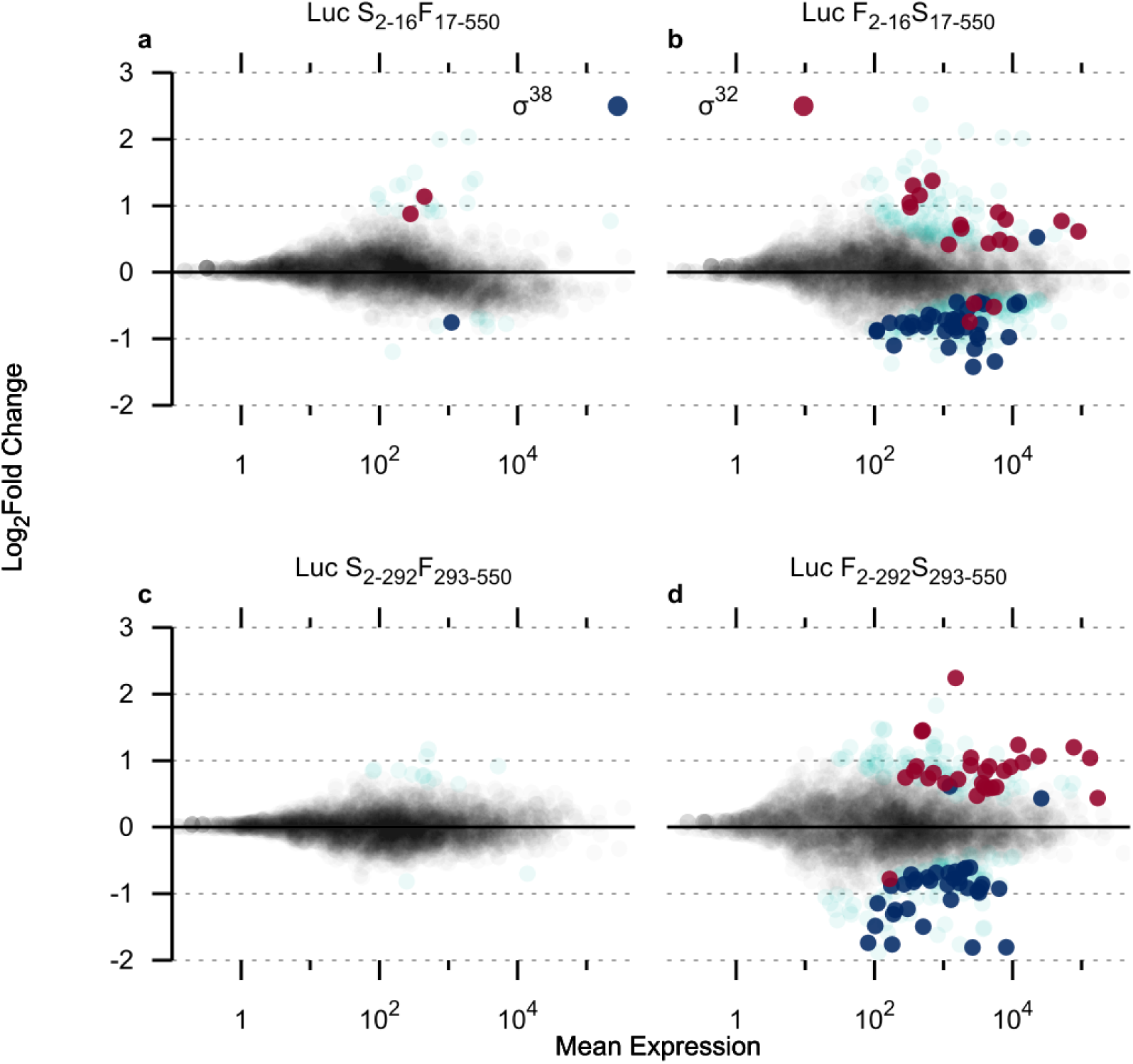
Quantification of ribosome-protected mRNA fragments mapped to mRNAs belonging to either the σ^32^ (red) or σ^38^ (blue) regulons. **a.** Luc S_2-16_F_17-550_ **b.** Luc F_2-16_S_2-550_ **c.** Luc S_2-292_F_293-550_ **d.** Luc F_2-292_S_293-550_. Red and blue circles: σ^32^ regulon and σ^38^ regulon, significant (p_adj_ < 0.10). Light blue circles: significant, other regulons. Grey circles: not significant.

**Figure S3:**
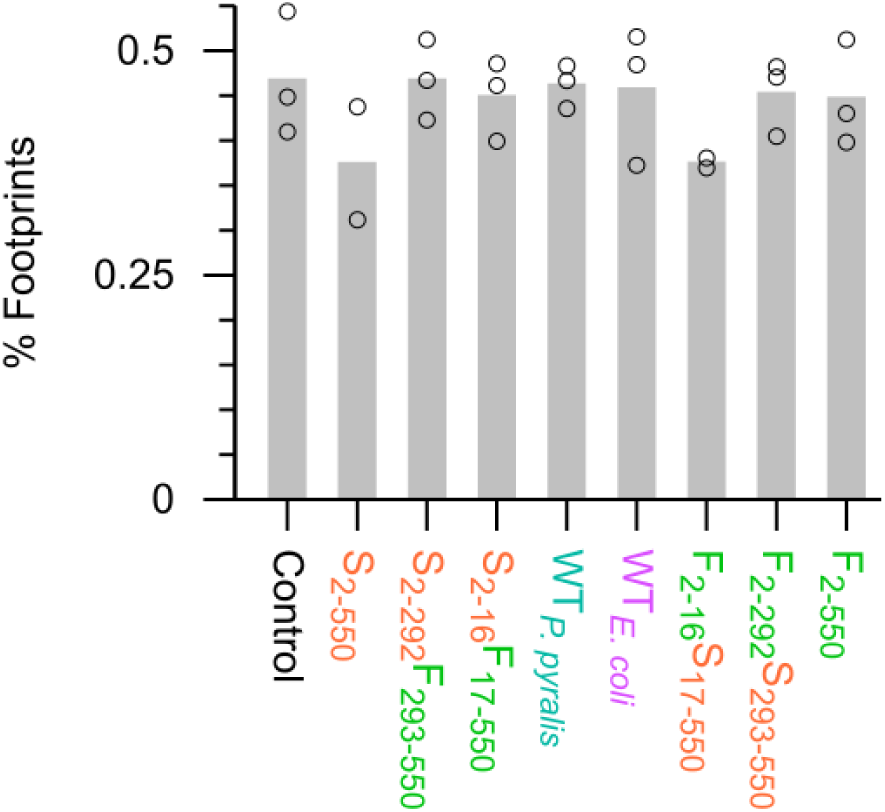
Quantification of ribosome-protected mRNA fragments mapped to the *tig* (trigger factor) mRNA in *E. coli* strains containing different synonymously re-coded *Luc* mRNAs. Circles denote experimental replicates (n = 2 or 3).

**Figure S4:**
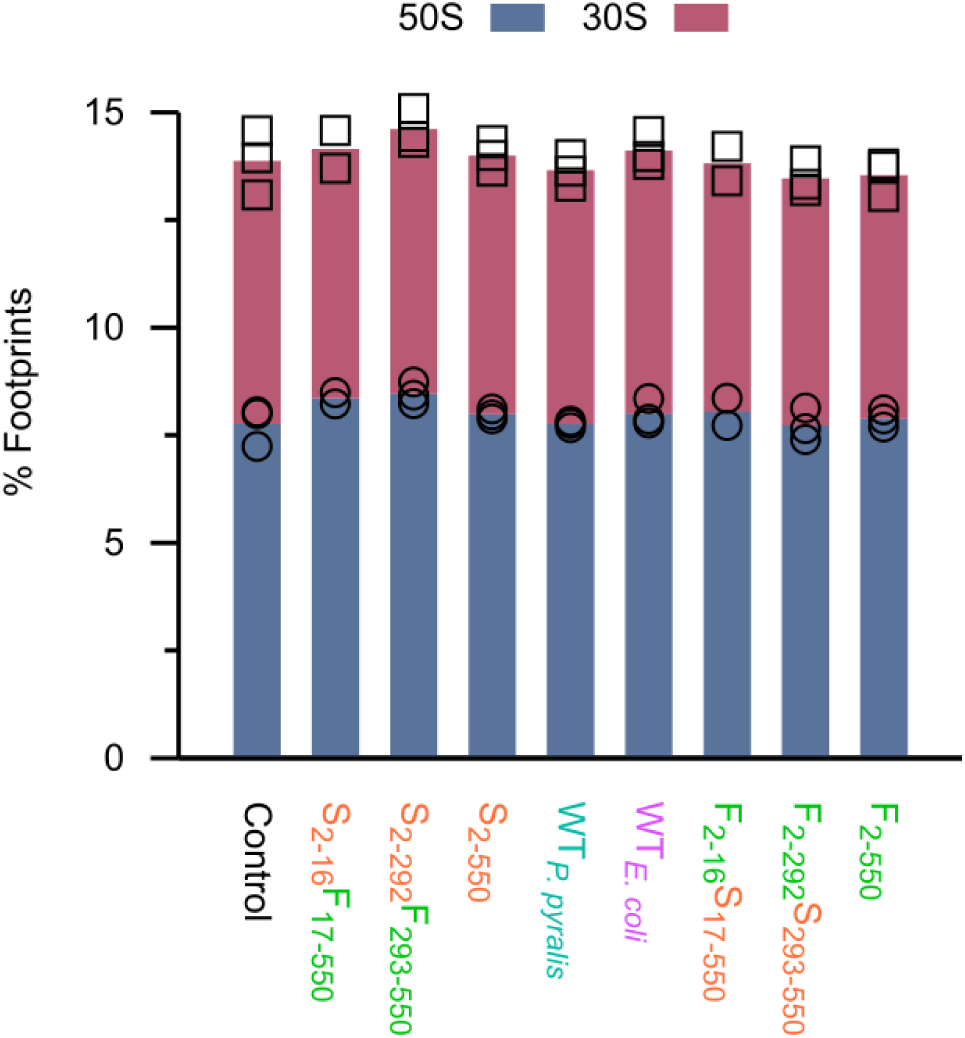
Quantification of ribosome-protected mRNA fragments mapped to the ribosomal proteins (50S subunit: blue; 30S subunit: red). Symbols denote experimental replicates (n = 2 or 3).

**Figure S5:**
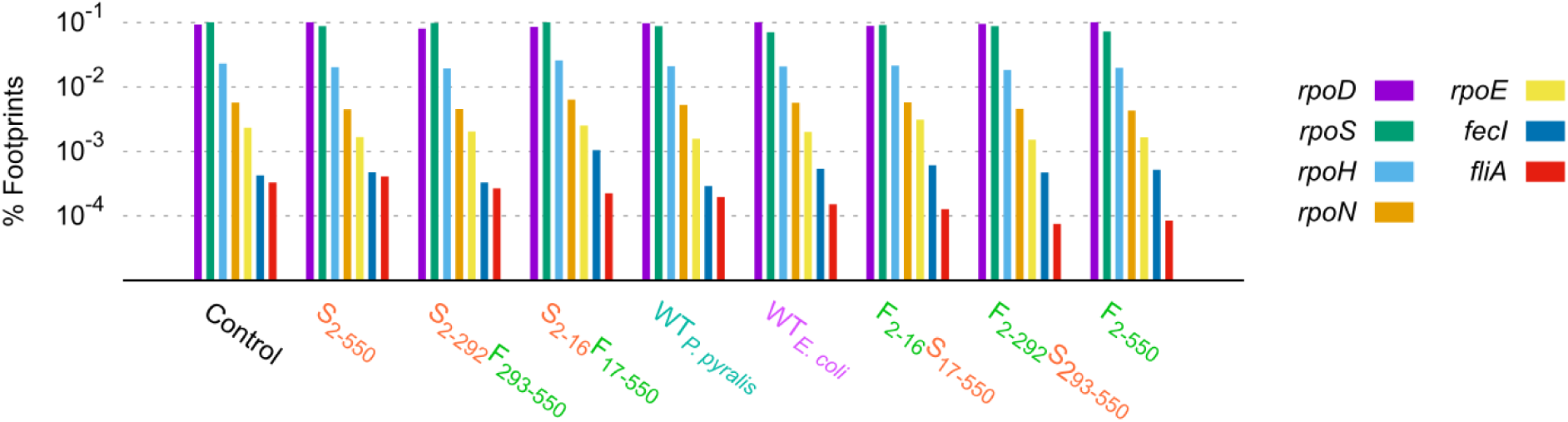
Quantification of ribosome-protected mRNA fragments mapped to the sigma factor mRNAs in *E. coli* strains containing different synonymously re-coded Luc mRNAs.

**Figure S6:**
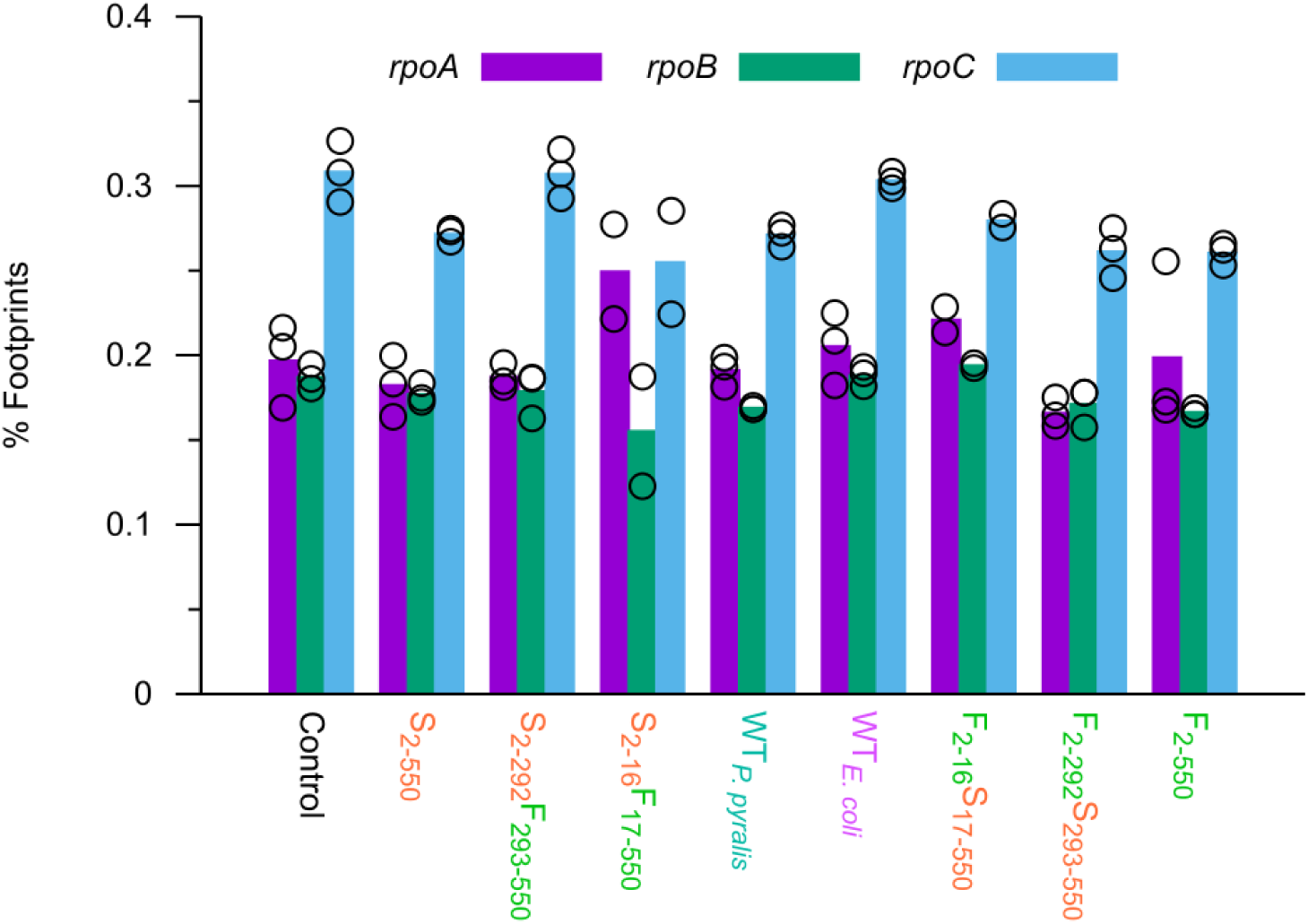
Quantification of ribosome-protected mRNA fragments mapped to the RNA polymerase subunits mRNAs in *E. coli* strains containing different synonymously re-coded Luc mRNAs. Symbols: replicates (n = 2 or 3).

## Methods

### Contact for Reagent & Resource Sharing

All sequencing data have been made available at the Gene Expression Omnibus (accession number GSE104303). Further information and requests for resources and reagents should be directed to and will be fulfilled by the Lead Contacts, Andrew T. Martens or Vincent J. Hilser. Code and procedures to process high throughput sequencing datasets was previously published by the authors and is available on GitHub (Martens et al., 2015).

### Experimental Model & Subject Details

#### Bacterial cell cultures

For all physiological experiments (ribosome profiling, mRNA sequencing, enzyme assays, immunoblotting) MOPS buffered medium (Neidhardt et al., 1974) plates (0.6% glycerol, 0.2% casamino acids, 100 μg/ml ampicillin) were streaked with *E. coli* MG1655 (Yale CGSC # 6300; F-, λ-, rph-1) cells bearing one of the pACYC177-derived plasmids and grown at 37°C (**Table S1**). Glycerol and casamino acids, rather than glucose, were added to minimize catabolite repression while avoiding amino acid starvation. Overnight cultures were prepared by inoculating 2.5 ml of liquid MOPS buffered medium, with 1 mM IPTG, from a plate, and grown overnight, shaking at 37°C at 250 RPM. Under these conditions, cells typically grow with a doubling time of 50 min. Other experiments, such as those necessary for the assembly of the pACYC177 plasmids (cloning), were performed using *E. coli* NEB 5-alpha cells grown in LB medium.

### Experimental Method Details

#### Preparation and transformation of competent cells

Transformations were carried out as previously described (Renzette, 2011), with minor changes. *E. coli* MG1655 cells were grown overnight in LB medium at 37°C, shaking at 250 RPM. The overnight culture was then diluted to an OD_600_ of 0.05 and grown until it reached an OD_600_ of 0.30. The culture was transferred to a 50 ml Falcon tube and then placed in an ice water bath in a 4°C cold room for 20 min. The culture was then centrifuged in a pre-chilled (4°C) Sorvall RC 5B Plus for 10 min at 3,000 RPM. After pouring off the supernatant, the cellular pellet was resuspended in 7.5 ml Transformation Buffer I (60 mM potassium acetate, 80 mM MnCl_2_ (added fresh), 100 mM KCl, 10 mM CaCl_2_, 1% glycerol) by gently swirling in the cold room ice water bath, whereupon it was left to incubate for 2 h. The chilled cells were re-centrifuged (10 min, 4°C, 3,000 RPM) and resuspended in 1 ml Transformation Buffer II (10 mM MOPS, 10 mM KCl, 100 mM CaCl_2_, 1% glycerol). 100 μl cellular aliquots were transferred to pre-chilled 1.5 ml Eppendorf microcentrifuge tubes and kept at −80°C until they were needed.

Competent cells were transformed with plasmid using the heat shock method. 1 to 5 μl of miniprepped plasmid (approximately 100 ng) were added to a single aliquot of cells kept on ice. After 30 min, the cells were placed in a 42°C water bath for exactly 30 s, and then placed again on ice for 2 min. After addition of 900 μl of SOC medium, the cells were incubated for 1 h at 37°C shaking at 250 RPM. The cells (between 100 and 400 μl) were then delicately spread on slightly dried LB ampicillin plates using a sterile, bent glass Pasteur pipette and allowed to grow overnight at 37°C.

#### Re-coding *Luc* mRNA using the translation index

A translation index was previously developed to rank synonymous codons according to WC/wobble and tRNA gene copy number parameters, and *Luc* mRNA variants were designed using this translation index (Spencer et al., 2012), what we refer to as Luc F_2-550_, Luc S_2-550_, and Luc WT*_E. coli_*. The unmodified wild-type mRNA (Luc WT*_P. pyralis_*) was also analyzed. Translation indices were compared by looking up each value at the corresponding position within the *Luc* coding sequence. These alleles all end with a peroxisome-inactivation sequence, a linker, a c-myc tag and a His_6_ tag, which were not re-coded.

#### Assembly of low-copy *Luc* vectors

To express *Luc* with high, robust levels of transcription, while simultaneously keeping total protein expression levels low enough to avoid overexpression artifacts, *Luc* alleles were cloned into the medium-copy number pACYC177 (Chang and Cohen, 1978) plasmid downstream of the pLac promoter and ribosome binding site sequences. PCR products were derived from vectors bearing synonymously re-coded *Luc* genes, which were previously generated (Spencer et al., 2012) (provided to us by J. Barral), and from the *E. coli* MG1655 chromosome; these were combined using the Gibson assembly method (Gibson et al., 2009) (**Table S3**). The *Luc* gene, including the transcriptional terminators from the pBAD vectors, was inserted on the strand opposite the one encoding the two antibiotic resistance genes to prevent transcriptional read-through. PCR products were prepared using the oligonucleotides listed in **Table S2**. Vector pACYC177 was digested with restriction enzyme BamHI-HF (New England Biolabs cat. no. R3136S), and the assemblies were performed as instructed by the kit (New England Biolabs cat. no. E2611).

#### Polymerase chain reactions

Amplification of DNA by PCR was performed using Q5 (New England Biolabs cat. no. M0491) or Phusion (New England Biolabs cat. no. E0553) polymerase (200 μM dNTP, 10 μM forward primer, 10 μM reverse primer, 1-50 ng DNA, 0.02 U/μl enzyme) in 50 μl reactions as directed by **Tables S2 & S3**, using a Veriti 96-well thermocycler (Applied Biosystems). *E. coli* MG1655 chromosomal DNA was prepared using the GeneJet Genomic DNA Purification Kit (ThermoFisher cat. no. K0721).

#### Ribosome profiling

200 ml of MOPS buffered medium, in pre-warmed 2 L flasks, were inoculated by diluting the overnight cultures more than one thousandfold and set to shake at 37°C in an Innova 40 incubator shaker (New Brunswick Scientific) at 200 RPM. Once the cultures reached an OD_600_ of 0.40 - 0.65 (measured using a Shimadzu UV-1800 spectrophotometer; 1 OD_600_ = 6 x 10^8^ cells / mL) they were filtered through a pre-warmed 0.22 μm nitrocellulose filter paper and 90 mm glass filtration system (Kontes) with vacuum (Whatman cat. no. 7182-009); the filtration took about 1 min. The apparatus was rapidly disassembled, and the cells were immediately scraped from the filter paper, using a pre-warmed scoopula, and submerged into a 50 ml Falcon tube (Corning cat. no. 352070) filled with liquid nitrogen (LN2). The cells were then transferred to a mortar immersed in, and containing, LN_2_ and 600 μl of cell lysis buffer (10 mM MgCl_2_, 100 mM NH_4_Cl, 20 mM Tris-HCl pH 8.0, 0.1% Nonidet P40, 0.4% Triton X-100, 500 μg/ml chloramphenicol) were dripped in. The cells were pulverized with the chilled mortar and pestle, whereupon the LN_2_ was allowed to boil off. The remaining powder was scraped with a chilled spatula into a chilled 1.5 ml Eppendorf tube and stored at −80°C overnight.

Cell lysates were centrifuged 15 min at 14,000 RPM and 4°C, and the clarified supernatant was transferred to a 1.5 ml Eppendorf tube. Between 250 μl and 500 μl of supernatant were then treated with 2.0 μl of micrococcal nuclease (Life Technologies cat. no. EN0181) and 5 mM CaCl_2_ and incubated 1 h while gently rotating at room temperature. The reactions were quenched using 6 mM EGTA and then transferred to 3.5 ml thickwall polycarbonate ultracentrifuge tubes (Beckman Coulter cat. no. 349622). The samples were underlaid with 2.0 ml of 1 M sucrose and centrifuged 6.5 h on a Beckman XL-80K ultracentrifuge at 55,000 RPM and 4°C in a SW55 rotor. Following removal of the liquid, the invisible pellets were re-suspended with 700 μl of Qiazol reagent, and RNA was extracted as directed by the miRNEasy kit (Qiagen cat. no. 217004).

Samples were denatured at 70°C and then subjected to electrophoresis on a 15% polyacrylamide TBE-urea gel 65 min at 200 V. The gel was then stained using SYBR gold and imaged with a blue light transilluminator. Gel slices were taken from the regions corresponding to lengths between 20 to 45 nt, as indicated by 10 bp ladder (Invitrogen cat. no. 10821015) and RNA size markers (5′-AUGUACACGGAGUCGAGCUCAACCCGCAACGCGA-(Phos)-3′ and 5′-AUGUACACGGAGUCGACCCAACGCGA-(Phos)-3′). The samples were then prepared as directed in Ingolia *et al*. ^(Ingolia et al., 2012)^, with modifications: ProtoScript II reverse transcriptase (New England Biolabs cat no. M0368) was used for reverse transcription, and different biotinylated oligonucleotides were used for the removal of *E. coli* rRNA sequences (Li et al., 2014). The library was sequenced on an Illumina HiSeq 2500 by the Johns Hopkins University Genetic Resources Core Facility. Each bacterial strain was processed in duplicate or triplicate from distinct biological samples on different days (**Table S1**).

#### Immunoblotting

2.5 mL of MOPS minimal medium (0.6% glycerol, 0.2% casamino acids, 100 μg/ml ampicillin (Neidhardt et al., 1974) were inoculated with an *E. coli* colony from a MOPS buffered medium plate and grown overnight, shaking 250 RPM at 37°C. Fresh 2.5 mL cultures were inoculated the following day to an OD_600_ of 0.01 and grown 6 hours (∼7 doublings) to an OD_600_ of ∼1.0 and then placed on ice. 1 mL aliquots were transferred to chilled 1.5 mL Eppendorf tubes and centrifuged 5 min at 14,000 RPM at 4°C. Using OD_600_ readings of the remaining cultures, the cellular pellets were resuspended in Laemmli sample buffer (50 μl / OD_600_), and the cellular suspensions were boiled 10 min at 100°C and centrifuged 2 min at 14,000 RPM at room temperature. 2.0 μl Amersham ECL Rainbow Marker (GE Health Care cat. no. RPN800E) and 1 μl cell lysate aliquots were subjected to gel electrophoresis by 10% SDS PAGE at 100 V for 80 min.

After equilibrating the gel in tris glycine buffer (25 mM Tris, 192 mM glycine, 15% methanol) for 30 min, transfer was executed using Immun-Blot LF PVDF membrane (Bio-Rad cat. no. 162-0261) in ice-cold tris glycine for 40 min at 100 V. After transfer, the membrane was allowed to dry 1 h. After recharging the membrane in methanol for 1 min, the membrane was rinsed in deionized water and washed for 2 min in TBS buffer (50 mM Tris, 150 mM NaCl, pH 7.6). The membrane was blocked in Odyssey Blocking Buffer (LI-COR cat. no. 927-50003) for 1 h at room temperature, then rinsed for 5 min in TBS-T (TBS, 0.1% Tween-20) three times. The primary antibodies (mouse anti-myc primary antibody (Cell-Signaling Technologies cat. no. 2276) for Luc detection and rabbit anti-β-galactosidase primary antibody (ThermoFisher cat. no. A-11132) were then diluted 1:20000 into Odyssey blocking buffer and 0.2% Tween-20, and allowed to incubate overnight at 4°C with gentle mixing. The membrane was then washed for 5 min in TBS-T three times. The secondary antibodies (goat anti-mouse IRDye 800CW secondary antibody (LI-COR cat. no. 925-32210) and goat anti-rabbit IRDye 680RD secondary antibody (LI-COR cat. no. 925-68071) were diluted 1:20000 fold into Odyssey blocking buffer and 0.2% Tween-20 and 0.01% SDS for 1 hour at RT, covered with foil. The membrane was then washed for 5 min in TBS-T three times, and a final wash in TBS for 5 min. After drying at RT for 1 h, the membrane was imaged using a LI-COR Odyssey Fc imaging system. Secondary antibody intensities were quantified using LI-COR Image Studio v. 5.2. The final ratio of Luc to β-galactosidase was calculated by dividing the total signals from the respective secondary antibodies.

#### Luciferase enzymatic assays

Luciferase activity was measured using the Dual-Light kit (ThermoFisher cat. no. T1004) with a Berthold TriStar2 plate-reader luminometer as follows. 2.5 mL MOPS buffered medium (0.6% glycerol, 0.2% casamino acids, 100 μg/ml ampicillin (Neidhardt et al., 1974) cultures were prepared from a minimal medium plate and grown overnight. These were then diluted into 2.0 mL of medium to an OD_600_ of 0.01 and grown to and OD_600_ of 1.0 at 37°C. The cultures tubes were then immediately placed on ice, and for each culture, three 100 μl aliquots of medium containing cells were transferred to 1.5 ml Eppendorf microcentrifuge tubes containing 100 μl of Tropix lysis solution. Cells were lysed by vortexing for 10 s. 10 μl of lysate were added to 25 μl of Buffer A (with luciferin) in a non-treated Nunc F96 MicroWell white polystyrene 96-well plate (ThermoFisher cat. no. 236105). After a single, initial 10 s shake step (diameter 1), Luciferase activity was sequentially stimulated by injection of 100 μl of Buffer B (speed 5), mixed for 0.2 s (diameter 5), and then, after a 2 s delay, measured continuously for 15 s with 0.05 s intervals. To reduce background signal, no galacton+ substrate was added to Buffer B. Measurements were performed at room temperature (∼23°C). The luminescence was then averaged over the 15 s duration, and the average blank value was subtracted from the average luminescence. After correcting for cell density (dividing by the OD_600_), 95% confidence intervals of the mean were calculated for the replicates (n = 3).

### Quantification & Statistical Analysis

#### Illumina sequencing data

FASTQ files were processed in a number of steps, leading to the assignment of integer counts of footprints per gene. First, the reads were trimmed by removing the adapter sequence, using fastx_clipper 0.0.13.2. Next, rRNA and tRNA sequences were filtered using Bowtie2 2.2.9 (Langmead and Salzberg, 2012). The remaining reads were mapped to the *E. coli* MG1655 genome (NC_00913.3) and the appropriate pACYC177 vector sequence using Bowtie2, followed by a procedure, written in Perl, which corrects sequencing errors and merges identical reads (Martens et al., 2015). Using the list of proteins and their corresponding genomic map points from PTT files, each gene was scanned for mapped reads and the totals were recorded to a new file. Reads mapped to *tufA* and *tufB* were merged, and an additional scan was performed to assign reads to *ssrA*. Differential gene expression analysis was performed using DeSeq2 1.12.3 (Love et al., 2014) and R 3.3.1.

#### *Luc* Ribosome profile density calculations

Densities along the re-coded *Luc* mRNA were compared as follows. Position-specific footprint densities were assigned from length-normalized 30-nucleotide, 3′-aligned (Martens et al., 2015) data, followed by base mean calculations from the replicates (**Table S1**) using DeSeq2 (Love et al., 2014), which uses a weighted averaging procedure. This procedure was used to not bias the average ribosome profile towards replicate datasets with higher sequencing depths. To compare the ribosome density along the mRNA, the profiles were window averaged (window size 151 nt), and the data were then normalized from 0 to 1, with 0 representing the true zero, and 1 representing the maximum value.

#### Differential ribosome allocation

Differential ribosome allocation was analyzed using DeSeq2 (Love et al., 2014). Integer numbers of sequencing reads, 1 per ribosome footprint, were assigned to either chromosomal *E. coli* genes or to genes encoded on the low copy pACYC177 vectors bearing *bla* (*ampR*), *kanR* and *Luc* genes. Since experiments on strains pACYC177 Control, Luc F_2-550_, Luc S_2-550_, LucWT*_E. coli_* and LucWT*_P. pyralis_* were done in parallel, these were analyzed both by *Luc* expression category as well as by replicate number, with pACYC177 datasets serving as the reference strain. The other experiments were only analyzed in terms of the *Luc* expression category, also with pACYC177 datasets serving as the reference strain. Adjusted p-values (p_adj_) were generated by DeSeq2 and used to determine statistically significant ribosome allocation (taking into account the number of replicates and sequencing depths) compared to control cells without *Luc*.

#### Correlations between immunoblotting and enzyme activity assays

Mean Luciferase enzymatic intensities were compared against the immunoblot Luc protein levels (normalized by β -galactosidase). The correlation coefficient was first calculated, using all datapoints, using the functionality built in the Gnuplot software. After using a linear regression, it was visually noted that strain S_2-16_F_17-550_ appeared to be an outlier; the correlation was re-calculated after omitting that single datapoint. Comparison of the residuals of the two fits, the two r^2^ values, and the two sums of the squares of the differences between the fits and the data confirmed S_2-16_F_17-550_ to be a statistical outlier.

#### Fitting the relationship between Luciferase production and σ^32^ activity

The following molecular mechanism was used to understand the relationship between *Luc* expression, using enzymatic activity as a proxy for the concentration of enzyme, and σ^32^ activity, defined as the percentage of sequencing reads mapped to genes regulated by σ^32^: To perform nonlinear least-squares fitting with error in both the independent and dependent variables, the standard errors of the mean were provided for both the ribosome profiling and luciferase activity measurements, and fits were performed using Orear’s effective variance method (to account for uncertainty in both dimensions) in Gnuplot v. 5.0.5.

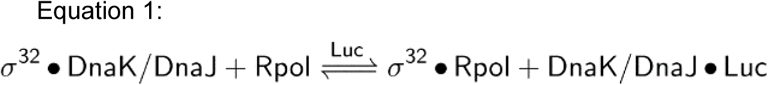

A single-site binding model, corresponding to the known molecular mechanism:

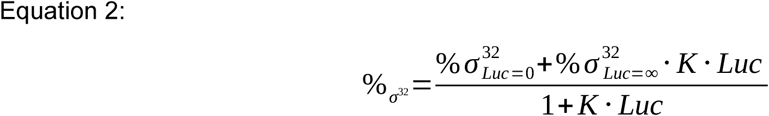

## Supplementary Tables

**Table S1:** *E. coli* MG1655 strains bearing plasmid pACYC177, number of ribosome profiling replicates and sequencing coverage.

| Insert | Name | Number of replicates | Luc sequence coverage (reads x 30 nt) (nucleotides) | Total mapped sequencing reads (all ORFs) |
| --- | --- | --- | --- | --- |
| None | Control | 3 | N/A | 15,602,867;<br>20,253,378;<br>16,550,616 |
| Slow Luc | Luc S<sub>2-550</sub> | 3 | 4,157,160;<br>6,763,380;<br>9,113,530 | 11,545,337;<br>14,060,269;<br>13,916,778 |
| Wild-Type Luc | Luc WT<sub>*P. pyralis*</sub> | 3 | 2,924,220;<br>17,965,350;<br>4,026,900 | 6,202,441;<br>23,283,404;<br>7,177,805 |
| Wild-Type Luc efficiency with *E. coli* codons | Luc WT<sub>*E. coli*</sub> | 3 | 6,455,670;<br>11,049,990;<br>6,776,310 | 8,921,131;<br>20,037,666;<br>13,084,734 |
| Fast Luc | Luc F<sub>2-550</sub> | 3 | 14,423,040;<br>23,850,030;<br>11,866,230 | 15,316,848;<br>25,456,674;<br>16,197,634 |
| Fast / Slow Luc | Luc F<sub>2-292</sub>S<sub>293-550</sub> | 3 | 13,548,060;<br>7,703,670;<br>8,596,230 | 8,781,449;<br>4,891,790;<br>5,501,023 |
| Slow / Fast Luc | Luc S<sub>2-292</sub>F<sub>293-550</sub> | 3 | 1,541,760;<br>207,210;<br>1,686,120 | 4,457,010;<br>937,606;<br>6,192,011 |
| Fast initial / Slow Luc | Luc F<sub>2-16</sub>S<sub>17-550</sub> | 2 | 5,389,440;<br>2,911,650 | 3,871,402;<br>2,288,255 |
| Slow initial / Fast Luc | Luc S<sub>2-16</sub>F<sub>17-550</sub> | 2 | 175,560;<br>524,850 | 848,480;<br>4,587,614 |

**Table S2:**
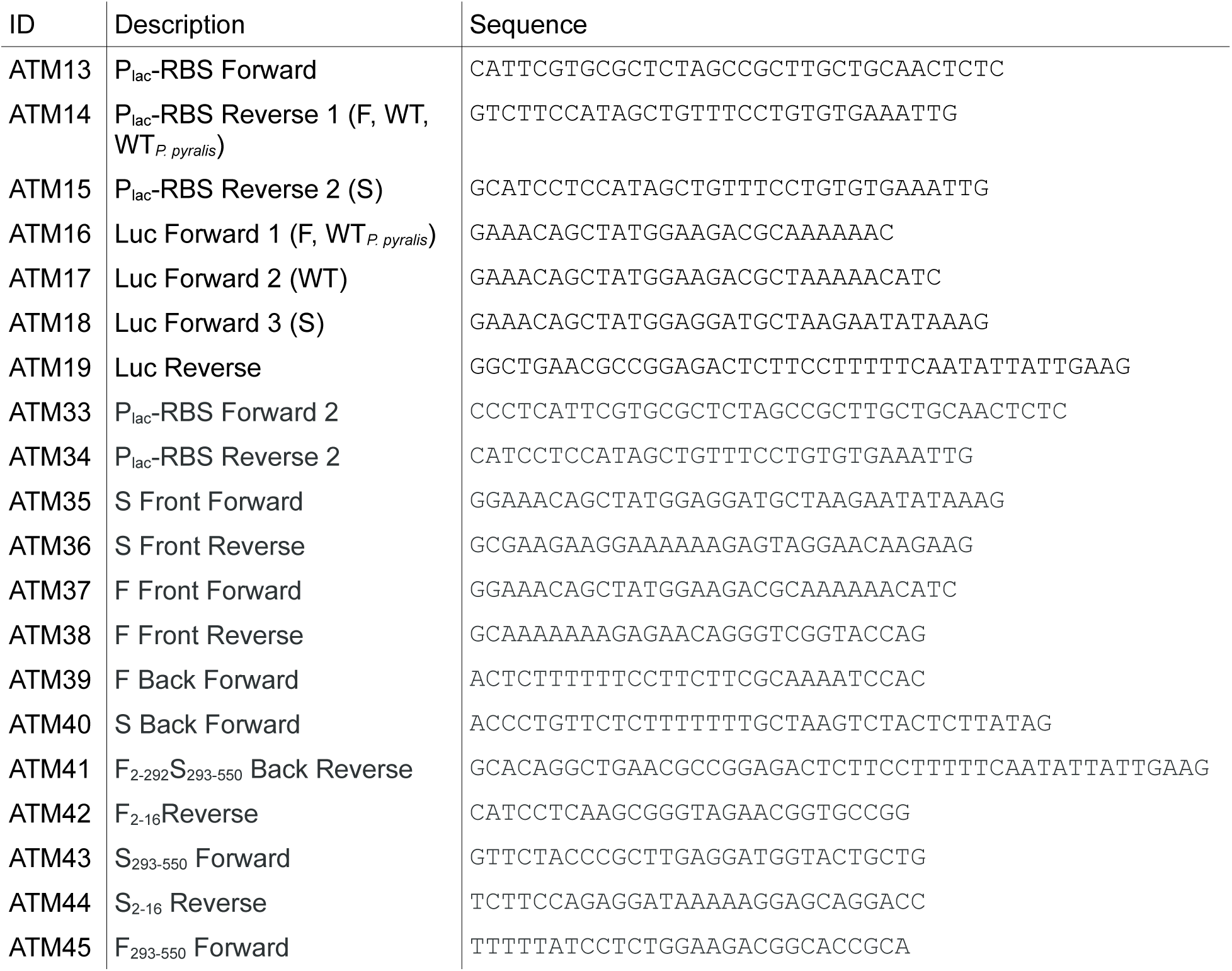
PCR Primers.

**Table S3:** Polymerase chain reactions (PCRs)

| DNA # | Forward Primer | Reverse Primer | Denaturation (initial) | Denaturation | Annealing | Extension | # Cycles | Template | Polymerase |
| --- | --- | --- | --- | --- | --- | --- | --- | --- | --- |
| 1 | ATM13 | ATM14 | 98°C/30s | 98°C/10s | 64.4°C/25s | 72°C/8s | 35 | *E. coli* MG1655 | Q5 |
| 2 | ATM13 | ATM15 | 98°C/30s | 98°C/10s | 64.4°C/25s | 72°C/8s | 35 | *E. coli* MG1655 | Q5 |
| 3 | ATM16 | ATM19 | 98°C/30s | 98°C/10s | 59°C/25s | 72°C/1m30s | 35 | pBAD18/LucF<sub>2-550</sub> | Q5 |
| 4 | ATM16 | ATM19 | 98°C/30s | 98°C/10s | 59°C/25s | 72°C/1m30s | 35 | pBAD18/LucWT<sub>*P. pyralis*</sub> | Q5 |
| 5 | ATM18 | ATM19 | 98°C/30s | 98°C/10s | 59°C/25s | 72°C/1m30s | 35 | pBAD18/LucS<sub>2-550</sub> | Q5 |
| 6 | ATM17 | ATM19 | 98°C/30s | 98°C/10s | 59°C/30s | 72°C/1m30s | 35 | pBAD18/lucWT<sub>*E. coli*</sub> | Q5 |
| 7 | ATM33 | ATM34 | 98°C/30s | 98°C/10s | 64.4°C/25s | 72°C/8s | 35 | *E. coli* MG1655 | Q5 |
| 8 | ATM35 | ATM36 | 98°C/30s | 98°C/10s | 60.5°C/25s | 72°C/50s | 35 | pACYC177/LucS<sub>2-550</sub> | Q5 |
| 9 | ATM33 | ATM38 | 98°C/30s | 98°C/10s | 60°C/25s | 72°C/50s | 35 | pACYC177/LucF<sub>2-550</sub> | Q5 |
| 10 | ATM39 | ATM41 | 98°C/30s | 98°C/10s | 60°C/25s | 72°C/50s | 35 | pACYC177/LucF<sub>2-550</sub> | Q5 |
| 11 | ATM40 | ATM41 | 98°C/30s | 98°C/10s | 60°C/25s | 72°C/50s | 35 | pACYC177/LucS<sub>2-550</sub> | Q5 |
| 12 | ATM13 | ATM42 | 98°C/30s | 98°C/5s | 68°C/10s | 72°C/1m8s | 35 | pACYC177/LucF<sub>2-550</sub> | Q5 |
| 13 | ATM13 | ATM44 | 98°C/30s | 98°C/5s | 65°C/10s | 72°C/9s | 35 | pACYC177/LucS<sub>2-550</sub> | Q5 |
| 14 | ATM45 | ATM19 | 98°C/30s | 98°C/5s | 68°C/10s | 72°C/1m8s | 35 | pACYC177/LucF<sub>2-550</sub> | Q5 |
| 15 | ATM43 | ATM19 | 98°C/30s | 98°C/10s | 60°C/10s | 72°C/1m30s | 35 | pACYC177/LucS<sub>2-550</sub> | Phusion |

## Notes

https://www.ncbi.nlm.nih.gov/geo/query/acc.cgi?acc=GSE104303

### Competing Interest Statement

The authors have declared no competing interest.

